# Cryo-EM of PMEL Amyloids Reveals Pathogenic Mechanism of Pigment Dispersion Syndrome

**DOI:** 10.1101/2024.12.09.627633

**Authors:** Haruaki Yanagisawa, Harumi Arai, Hideyuki Miyazawa, Masahide Kikkawa, Toshiyuki Oda

**Affiliations:** Department of Cell Biology and Anatomy, Graduate School of Medicine, the University of Tokyo, 7-3-1 Hongo, Bunkyo-ku, Tokyo, 113-0033, Japan; Department of Anatomy and Structural Biology, Graduate School of Medicine, University of Yamanashi, 1110 Shimokato, Chuo, Yamanashi, 409-3898, Japan

## Abstract

PMEL amyloids provide a vital scaffold for melanin deposition in melanosomes, playing a central role in pigmentation. Despite their importance, the high-resolution structure of PMEL amyloids has remained elusive. Here, we determined near-atomic resolution structures of wild-type PMEL amyloids using cryo-electron microscopy, revealing two distinct polymorphic forms with unique structural features. We further examined the pathogenic G175S mutation linked to pigment dispersion syndrome (PDS). Structural analysis showed that the G175S mutation introduces an additional hydrogen bond, stabilizing a novel fibril conformation. In vitro assays demonstrated a fourfold increase in polymerization efficiency for the G175S mutant compared to the wild-type. This enhanced polymerization correlated with a ∼70% increase in secreted amyloids in G175S-expressing cells without detectable changes in melanosome morphology or number. These findings suggest that the G175S mutation promotes amyloidogenesis within melanosomes, increasing amyloid load and contributing to PDS pathophysiology. This study provides insights into the molecular basis of PMEL amyloid formation in both physiological and pathological contexts, offering new perspectives on their structural diversity and dysregulation in pigmentation disorders.

## Introduction

Amyloids are protein aggregates traditionally associated with neurodegenerative diseases, but they can also play essential roles in normal physiological processes ^1–3^. One example is PMEL (Pmel17/gp100), a pigment-cell-specific protein that forms amyloid fibrils in melanosomes to scaffold melanin deposition. These fibrils underscore the dual nature of amyloids as both pathological and functional entities ^4–6^.

PMEL amyloidogenesis occurs within specialized organelles called melanosomes, which progress through four distinct stages (I–IV) of maturation ^7^. PMEL fibrils form during stage II and are essential for the transition to stage III, where melanin deposition begins. The structural organization of these fibrils is integral to their function, yet their high-resolution structure has remained unresolved. While the core amyloid-forming (CAF) domain and repeat (RPT) domain have been implicated in amyloid formation, their precise structural contributions are not fully understood ^8–12^ ^13^ Notably, glycosylation of the RPT domain has been suggested to modulate fibril organization ^14–16^.

Mutations in the PMEL gene, such as Gly175Ser (G175S), are associated with pigment dispersion syndrome (PDS), a condition in which pigment granules are released into the anterior chamber of the eye. This aberrant pigment release can elevate intraocular pressure, increasing the risk of pigmentary glaucoma (PG) ^17^. Approximately 15–20% of PDS patients develop PG, which can lead to vision loss ^18^. Although the G175S mutation has been proposed to alter PMEL amyloid formation, its precise molecular and structural effects remain unclear ^19^.

Here, we present near-atomic resolution structures of native PMEL amyloids, revealing two polymorphic forms in wild-type fibrils and structural alterations caused by the G175S mutation. Our findings link accelerated amyloidogenesis in the G175S mutant to increased amyloid load in melanosomes, providing insights into the molecular underpinnings of PDS and functional amyloid biology.

## Results

### Cryo-EM Structure of Native PMEL Amyloids Reveals Two Polymorphs

Native PMEL amyloids were isolated from the human melanoma cell line HMV-II through deglycosylation, protease digestion, and sonication. Cryo-electron microscopy (cryo-EM) and 2D classification revealed two distinct fibrillar forms: thick and thin filaments, corresponding to two-protofilament and single-protofilament structures, respectively (Figure 1a and S1). High-resolution structures were obtained from two-protofilament fibrils as shown in Figure 1b. However, our attempts to resolve the single-protofilament structures were unsuccessful, possibly due to their inherent instability or disruption during the extraction and processing stages. This suggests that the single-protofilament forms may represent immature or structurally less stable amyloid species, which are challenging to analyze at high resolution under the conditions applied in this study.

**Figure 1:**
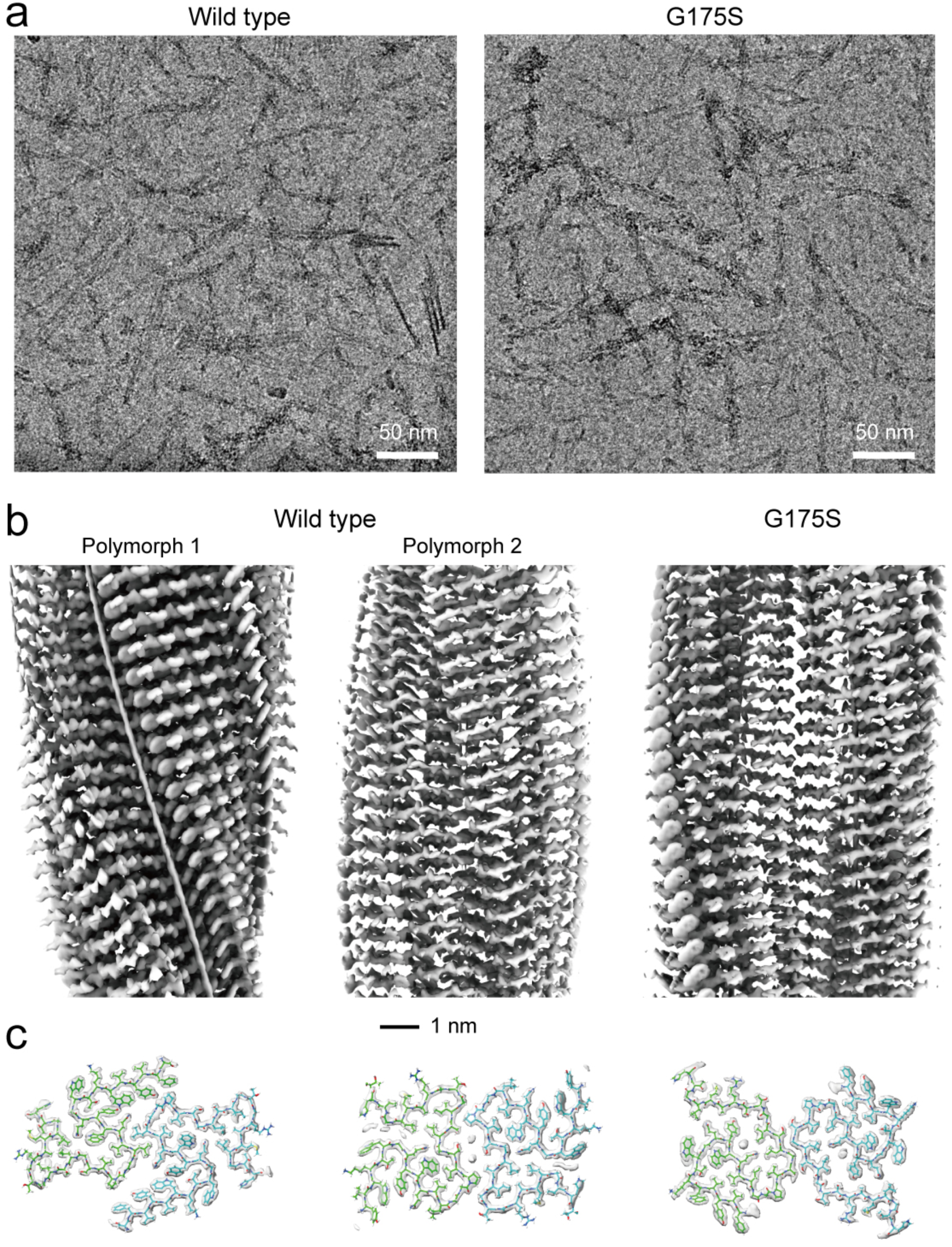
Cryo-EM structures of native PMEL amyloids. (a) Cryo-EM images of native PMEL amyloids extracted from the human melanoma cell line, showing both thick (two-protofilament) and thin (single-protofilament) fibrils (Figure S1, squares). (b) Side views of the reconstructed 3D maps of the two-protofilament fibrils, illustrating the overall helical structure of the amyloids. (c) Cross-sectional views of the reconstructed maps (gray), superimposed with models (green and cyan) of the CAF domain.

Two polymorphic forms of two-protofilament fibrils, termed Polymorph 1 and Polymorph 2, were identified in the native amyloids (Figure 1b and 1c). Polymorph 1 adopts a two-start helical architecture, characterized by a helical twist of 177.7° and a helical rise of 2.34 Å. In contrast, Polymorph 2 forms a one-start helical architecture with C2 symmetry, featuring a helical twist of -4.1° and a helical rise of 4.67 Å. Despite similar β-sheet configurations, the main chain morphology between the two polymorphs is markedly different (Figure 2a-c). Notably, Polymorph 2 encompasses a central cavity, which is notably absent in Polymorph 1, thereby rendering Polymorph 1 as a more densely packed structure. This central cavity may play a pivotal role in mediating fibril interactions with melanin precursors or other components intrinsic to the melanosome.

**Figure 2:**
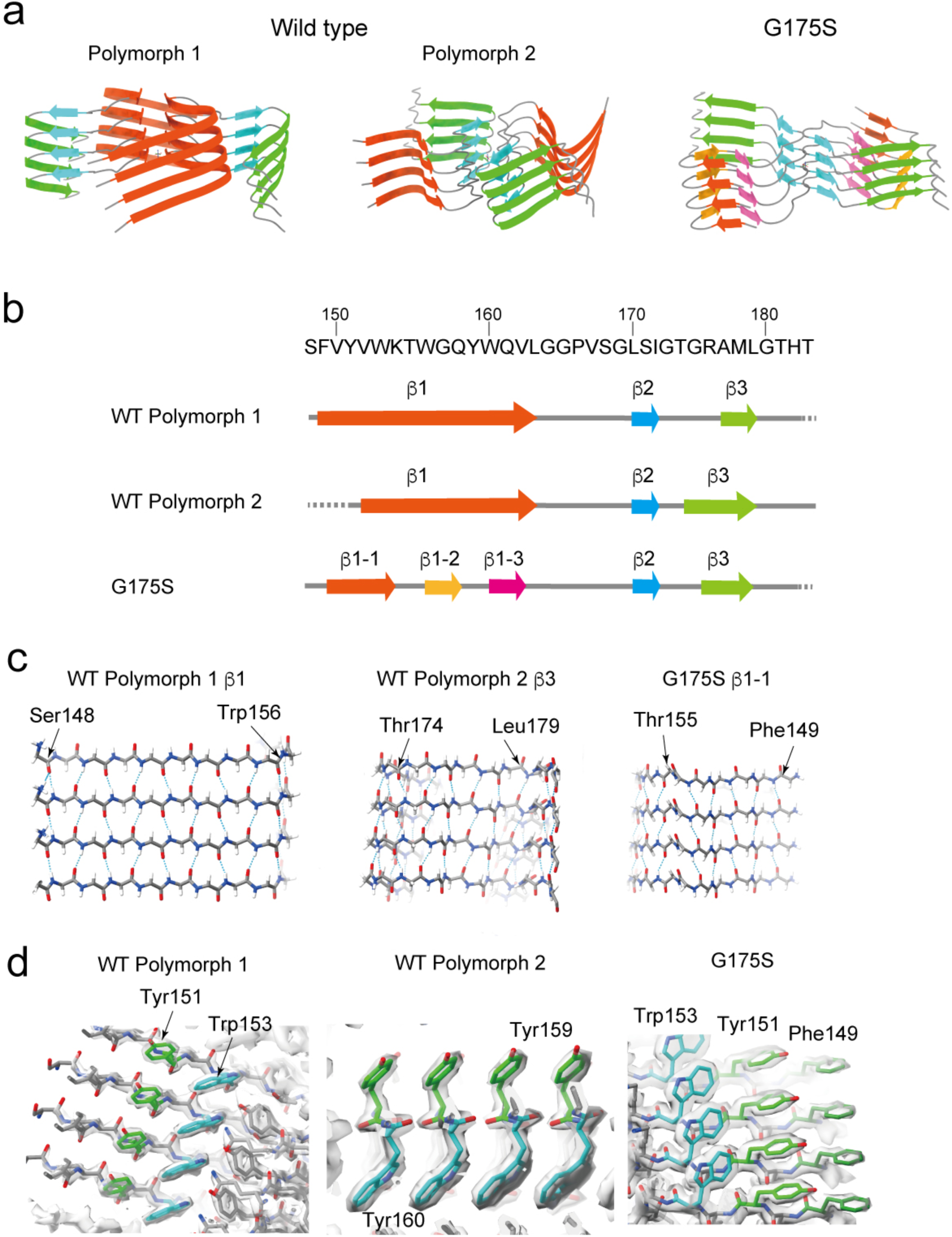
β-strand configuration of native PMEL amyloid. (a) Ribbon diagrams of native PMEL amyloid fibrils, illustrating the four-layered structure of β-strands in the fibril core. (b) Amino acid sequence of the N-terminal portion of the CAF domain, with the positions of β-strands (indicated by arrows) predicted using ModelAngelo. (c) Close-up views of the parallel cross-β sheets, highlighting the structural differences between wild-type and G175S mutant fibrils. In the G175S mutant, Thr155 separates β1-1 and β1-2, resulting in the division of the β-sheet. (d) Alignment of aromatic residues along the fibril axis, emphasizing the positioning of these bulky residues in both wild-type and G175S mutant fibrils.

Cryo-EM maps revealed that the side chains of aromatic residues, including tyrosine (Tyr), phenylalanine (Phe), and tryptophan (Trp), are aligned along the fibril surface. However, these residues do not form π-π stacking interactions, as their rings are not directly aligned. Instead, the alignment of hydrophobic bulky side chains likely stabilizes the amyloid structure and facilitates interactions with melanin during deposition (Figure 2d).

### Structural Alterations in G175S Mutant Amyloids

To investigate the effects of the G175S mutation on PMEL amyloids, we expressed G175S PMEL in HMV-II melanoma cells and isolated the resulting amyloids (Figure 1-2, left). Similar to the wild-type, cryo-EM revealed thick and thin filaments, with high-resolution reconstructions obtained exclusively from the thick filaments (Figure 3a and S2). The G175S amyloids exhibited a two-start helical architecture with a helical twist of 178.4° and a helical rise of 2.35 Å, similar to Polymorph 1 in wild-type amyloids. However, the β-sheet configuration and packing within the G175S fibrils were markedly altered.

**Figure 3.**
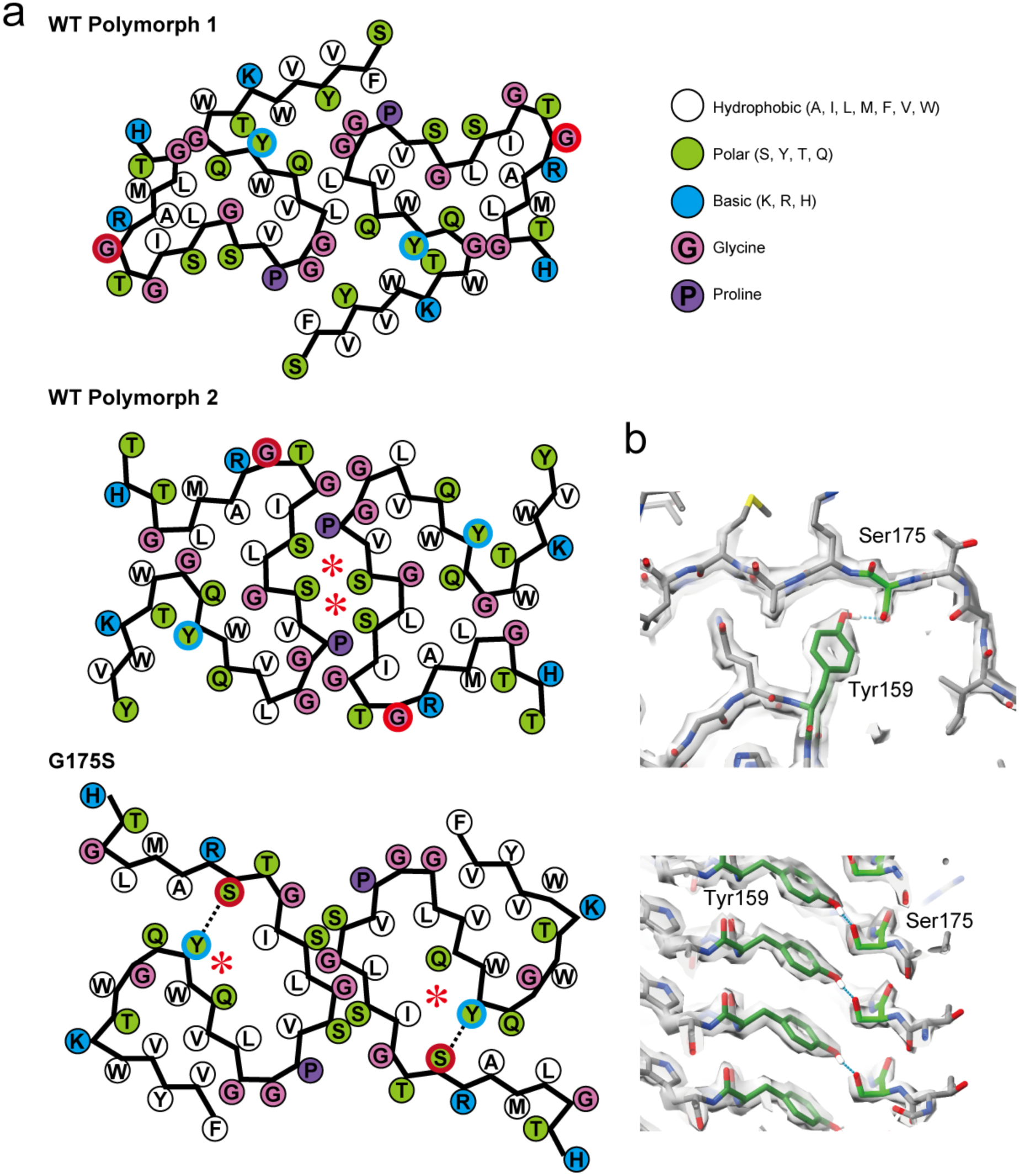
Additional hydrogen bond in G175S alters packing. (a) Packing schemes of one cross-sectional layer of the fibrils, illustrating the structural arrangement of residues in wild-type and G175S mutant fibrils. Asterisks (*) indicate the inner cavities present in Polymorph 2 and G175S. Blue and red circles highlight the positions of Tyr159 and Gly/Ser175, respectively. (b) Close-up view of the hydrogen bond between Tyr159 and Ser175 in the G175S mutant fibril, showing how this additional interaction alters the packing of the fibril compared to the wild-type.

In G175S amyloids, the first β-sheet (β1) is divided into three distinct segments (β1-1, β1-2, and β1-3), in contrast to the single continuous β1 observed in wild-type amyloids. β2 and β3 maintain similar configurations to those in the wild-type, but the overall packing density is reduced due to the division of β1 (Figure 2a-c). A notable structural feature in G175S amyloids is the formation of an additional hydrogen bond between Ser175 and Tyr159, which may contribute to the fragmented β1 configuration and potentially enhance fibril stability (Figure 3b).

### In Vitro Polymerization of Wild-Type and G175S PMEL CAF Domains

To further investigate the biochemical properties of PMEL amyloids, we expressed and purified the CAF domain (residues 148–223) of wild-type and G175S PMEL in *E. coli*. The purified proteins were subjected to *in vitro* polymerization assays, and the resulting amyloids were analyzed using cryo-EM and thioflavin T (ThT) fluorescence.

Cryo-EM analysis revealed that the wild-type CAF domain polymerized into fibrils structurally identical to Polymorph 1 of native PMEL amyloids, confirming that the *in vitro* polymerization recapitulates the native fibril structure. Similarly, the G175S CAF domain polymerized into fibrils indistinguishable from the G175S native amyloids, further validating that the extraction process from melanoma cells did not disrupt the amyloid structure (Figures 4a-c, S3-4). Notably, no one-start helical fibrils were observed in the in vitro polymerized samples, suggesting that thin fibrils represent immature intermediates or are disassembled during the extraction process.

**Figure 4:**
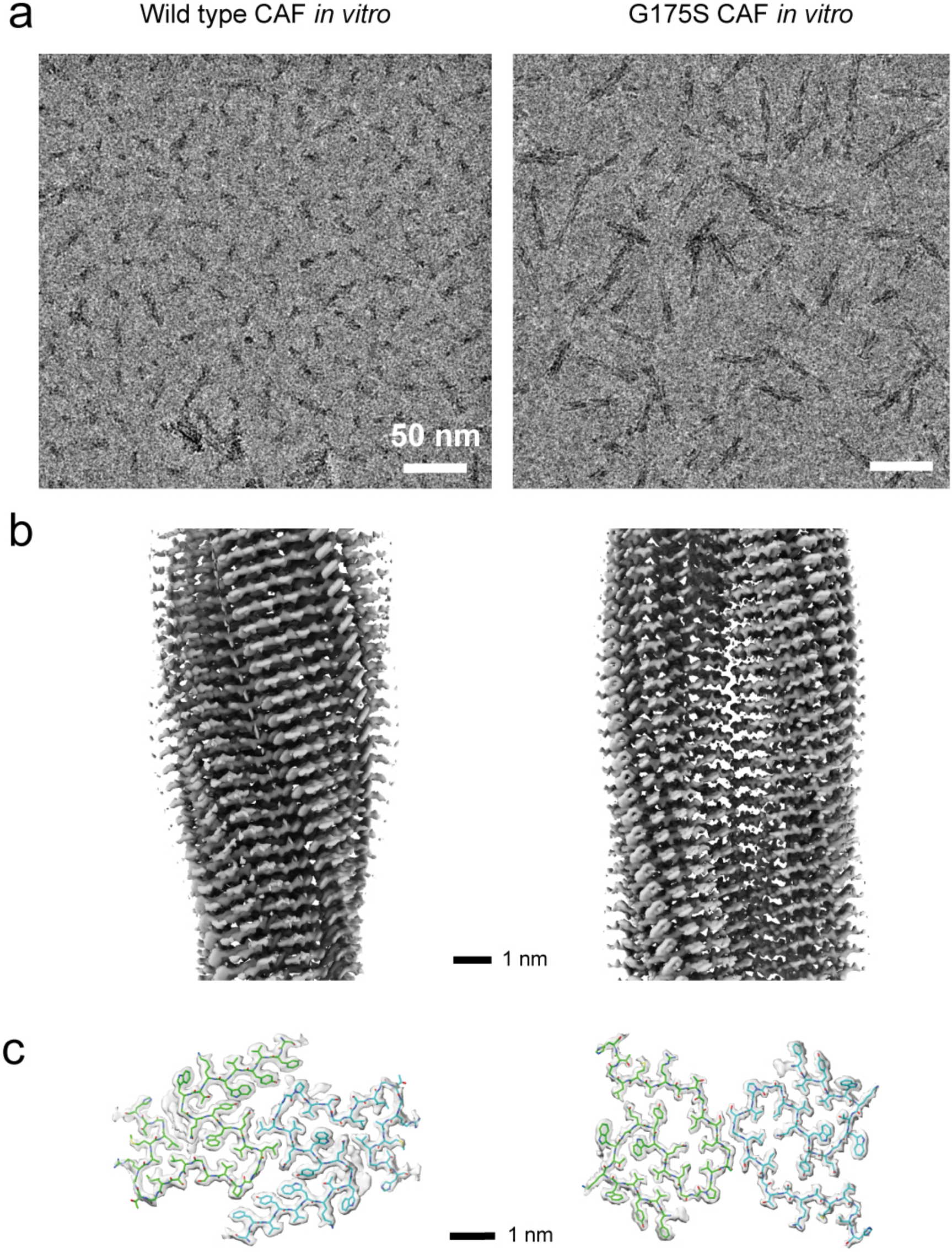
Cryo-EM of PMEL CAF domain fibrils polymerized *in vitro*. (a) Cryo-EM images of PMEL CAF domain fibrils polymerized *in vitro*, showing thick fibrils corresponding to two-protofilament structures. Wild-type fibrils are shorter than G175S ones. (b) Side views of the reconstructed 3D maps of the *in vitro* polymerized fibrils, highlighting the overall helical structure. (c) Cross-sectional views of the reconstructed maps (gray), superimposed with models (green and cyan) of the CAF domain.

### G175S mutation enhances amyloid formation in vitro and in vivo

Next, we performed ThT fluorescence assays to measure the amyloid-forming efficiency of wild-type and G175S CAF domains *in vitro* ^20^ (Figure 5a-b). The G175S mutant polymerized approximately four times faster than the wild-type, as shown by the rapid increase in ThT signal. Additionally, both G175S and wild-type polymerization curves plateaued, demonstrating that the final yield of polymerized amyloids is approximately four times greater for the G175S mutant than for the wild-type. The increased amyloid-forming efficiency of the G175S mutant is consistent with its structural alterations, particularly the possible stabilizing effect of the additional Ser175-Tyr159 hydrogen bond. The results from *in vitro* polymerization assays provide a mechanistic basis for the enhanced amyloidogenesis observed in G175S mutant cells. Despite the same amount of PMEL protein, the G175S mutation can increase the proportion of amyloid fibrils formed, thereby elevating the amyloid content within the melanosomes.

**Figure 5:**
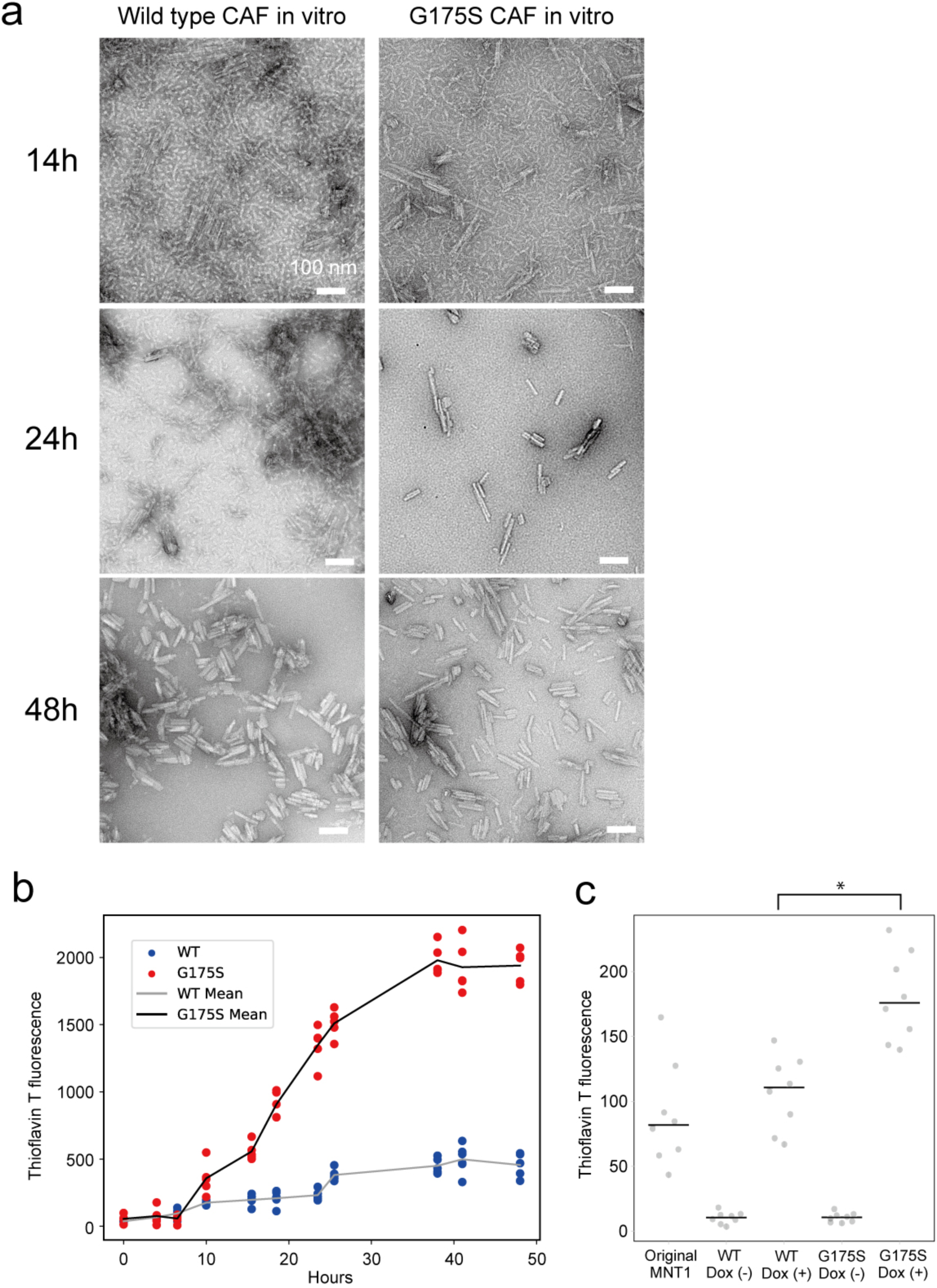
Enhanced amyloid-forming capacity of G175S CAF domain. (a) Negative-stain electron microscopy images of CAF domain fibrils polymerized *in vitro*. Thick amyloids are visible after 24 hours for wild-type and 14 hours for G175S. In the G175S mutant, even thicker amyloid bundles form at 24 hours, whereas wild-type fibrils only form comparable bundles after 48 hours. (b) ThT fluorescence assays quantifying polymerized amyloids. The plot shows the mean fluorescence values along with individual data points (N = 5), illustrating the faster and more efficient polymerization of the G175S mutant compared to the wild-type. A Mann-Whitney U test was used to compare the yield of polymerized amyloids between G175S and wild-type, showing a significant difference (*p* = 7.937e-3). (c) Thioflavin T fluorescence assays of secreted amyloids in cell culture supernatant. The plot shows the mean fluorescence values (lines) along with individual data points (N = 8). The fluorescence values were adjusted to reflect secreted amyloids from 1 × 10^7^ cells per data point. Original MNT1 refers to the amount of wild-type PMEL amyloids, while WT/G175S Dox (-) represent the knockout cell line without PMEL expression. WT/G175S Dox (+) show amyloid levels after 0.2 µg/ml doxycycline (Dox) induction of WT or G175S PMEL for 96 hours. The G175S Dox (+) condition resulted in approximately a 70% increase in amyloid secretion compared to WT Dox (+). A one-way ANOVA showed significant differences between groups (*p* = 7.216e-15), and Tukey’s HSD post-hoc test confirmed that the difference between WT Dox (+) and G175S Dox (+) was statistically significant (*p* = 3.362e-5) (asterisk).

### G175S mutation increases amyloid secretion

To evaluate the impact of the G175S mutation on amyloid secretion, we quantified secreted amyloids in the culture supernatant of MNT1 cells expressing wild-type or G175S PMEL using ThT fluorescence assays (Figure 5c). MNT1 cells expressing wild-type PMEL secreted amyloids at levels comparable to those of original MNT1 cells, consistent with the functional role of PMEL amyloids in pigmentation. In contrast, cells expressing the G175S mutant PMEL secreted significantly higher levels of amyloids—approximately 70% more than the wild-type. Notably, cells lacking PMEL expression showed negligible ThT fluorescence, confirming that the observed signal was specifically attributable to PMEL amyloids. Statistical analysis using one-way ANOVA revealed a significant difference among the groups (*p* = 7.216e-15). Tukey’s HSD post-hoc test confirmed that the difference between wild-type and G175S PMEL secretion was statistically significant (*p* = 3.36e-5). The increased secretion of amyloids in G175S cells aligns with the accelerated polymerization kinetics observed *in vitro*, suggesting that the mutation enhances amyloid formation within melanosomes.

### G175S mutation does not affect melanosomal structure in situ

To investigate whether the G175S mutation affects melanosome architecture, we employed cryo-focused ion beam scanning electron microscopy (cryo-FIB-SEM) and cryo-electron tomography to analyze melanosomes in MNT1 cells expressing wild-type or G175S PMEL (Figure 6a). Stage III melanosomes, characterized by fibrillar lamellae, were the primary focus of this analysis ^12, 21^.

**Figure 6:**
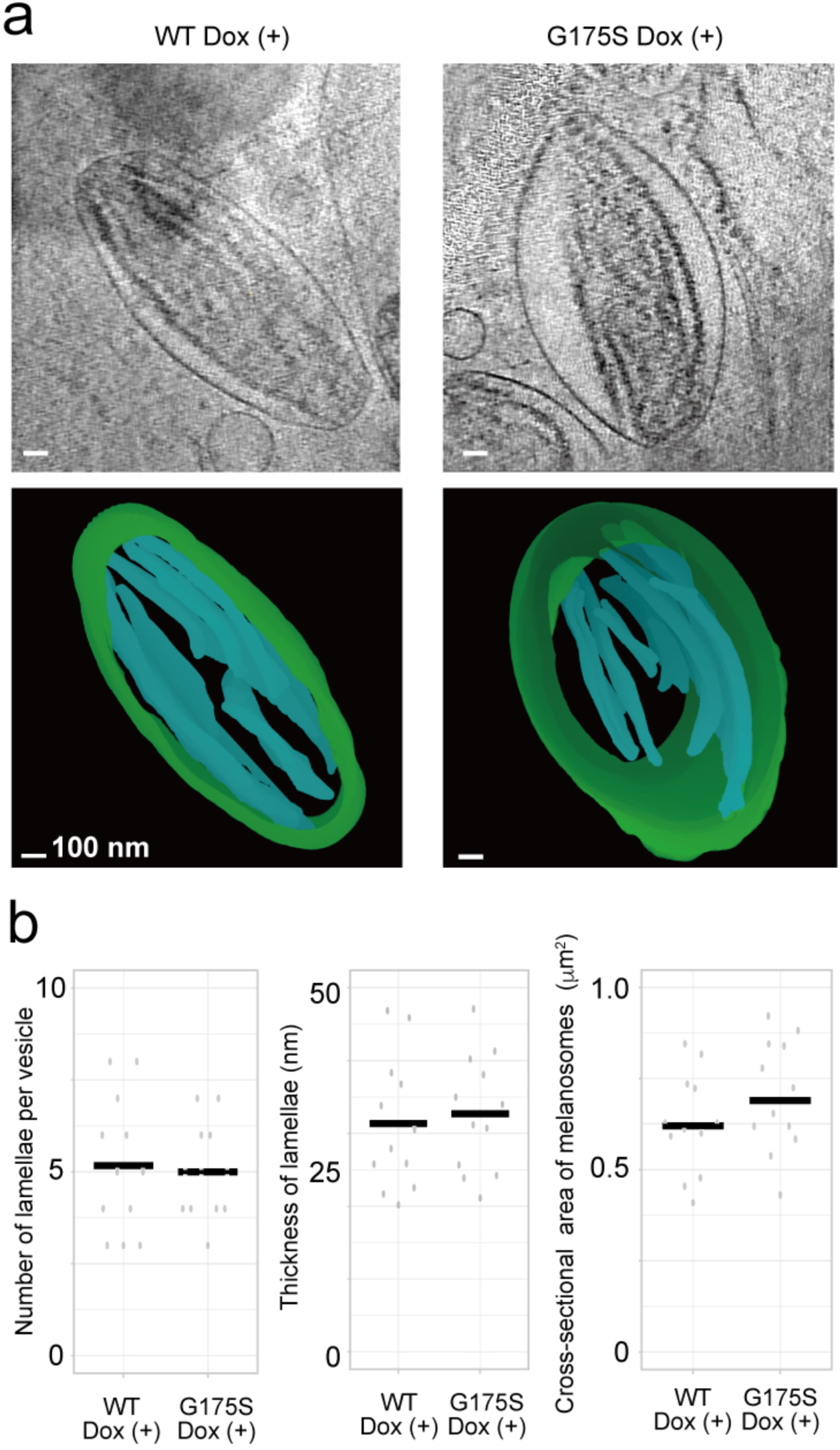
Structural analysis of melanosomes in MNT1 cells using cryo-FIB-SEM Tomography. (a) Cryo-tomographic slices (top panels) and 3D models (bottom panels) of stage III melanosomes from MNT1 cells expressing either WT or G175S PMEL, with Dox induction. Green represents the melanosomal membrane, and cyan indicates the fibrillar lamellae within the melanosome. Scale bars, 100 nm. (b) Quantification of structural parameters within melanosomes. The number of lamellae per melanosome, thickness of lamellae, and cross-sectional area of melanosomes were measured for WT Dox (+) and G175S Dox (+) cells. No significant differences were observed in the number of lamellae, lamellar thickness, or cross-sectional area of melanosomes between the two groups (N = 12). Individual data points are shown, with horizontal bars indicating the median values.

Tomographic slices revealed no discernible differences in the overall structural organization of melanosomes between wild-type and G175S mutant cells. Both cell types displayed lamellar structures, which were reconstructed into three-dimensional models to further examine the spatial arrangement of the fibrillar networks. Quantitative analysis demonstrated no significant differences in the number of lamellae per melanosome or the thickness of the lamellae between wild-type and G175S melanosomes (Figure 6b). Additionally, the cross-sectional area of melanosomes was consistent across the two groups.

These observations suggest that the G175S mutation, despite accelerating amyloidogenesis, does not alter the gross morphology of melanosomes. The increased secretion of amyloids observed in G175S mutant cells (Figure 5c) may, therefore, result from enhanced amyloid polymerization within the melanosome rather than structural changes that facilitate amyloid release.

### G175S affects the progression of melanosomal maturation stages

To determine whether the G175S mutation affects melanosome maturation, we quantified melanosome density and analyzed the distribution of melanosome stages (II, III, IV) using ultrathin-section electron microscopy of MNT1 cells expressing wild-type or G175S PMEL (Figure 7a) ^22, 23^. Melanosome density, calculated as the number of melanosomes per square micrometer of cytoplasm, showed no significant difference between wild-type and G175S mutant cells, indicating that the mutation does not alter melanosome abundance (Figure 7b, left panel).

**Figure 7.**
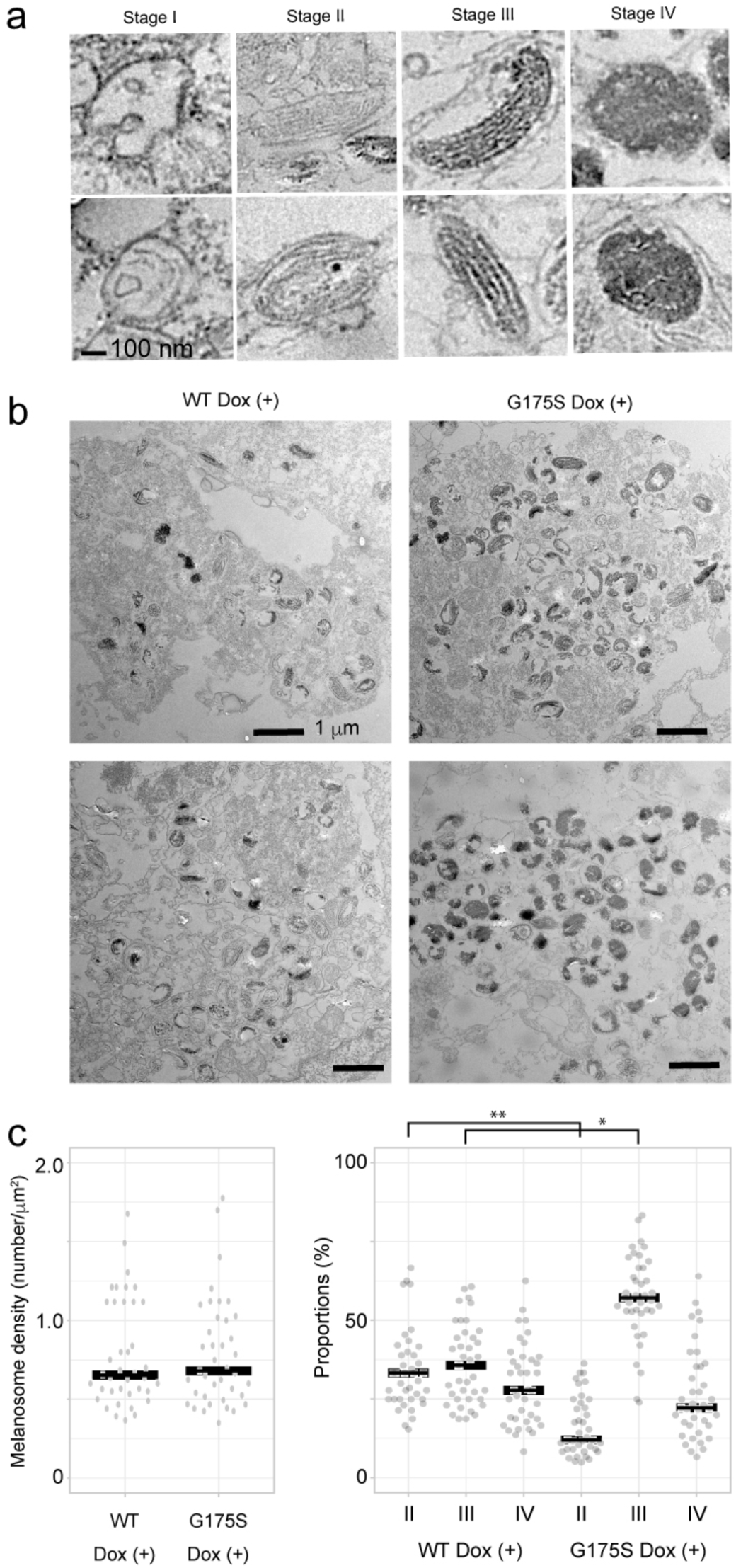
Melanosome morphology and stage distribution in MNT1 cells expressing wild-type or G175S PMEL. (a) Representative images of melanosomes in stages I–IV. Stage I: multi-vesicular bodies. Stage II: vesicles containing fibrillar lamellae without melanin deposition. Stage III: vesicles containing fibrillar lamellae with melanin deposition. Stage IV: vesicles with fully melanized, electron-dense granules. (b) Ultrathin-section electron microscopy images of MNT1 cells expressing wild-type (WT) or G175S PMEL with Dox induction. Melanosomes are visible as dark, electron-dense structures distributed throughout the cytoplasm. (c) Quantification of melanosomes. Left: Melanosome density (number per square micrometer of cytoplasm). Right: Percentage of melanosomes in stages II, III, and IV. Horizontal bars represent median values, and dots represent individual data points (N = 39 cells for each condition). A Wilcoxon signed-rank test revealed a significant increase in stage III melanosomes in G175S cells compared to WT cells (*p* = 2.575e-7, asterisk), while the densities of melanosomes did not significantly differ between conditions. Stage I melanosomes were not included in the quantification due to their low abundance and difficulty in identification.

Next, we assessed the proportion of melanosomes in stages II, III, and IV. The G175S mutation significantly increased the proportion of stage III melanosomes compared to wild-type, as determined by a Wilcoxon signed-rank test (*p* = 2.575e-7) (Figure 7b, right panel). Additionally, the proportion of stage II melanosomes was significantly decreased in G175S cells compared to WT (*p* = 1.568e-8, Figure 7b, double asterisk), while no significant differences were observed in the proportion of stage IV melanosomes (*p* = 0.3607). These results suggest that the G175S mutation accelerates the transition from stage II to stage III melanosomes without affecting the overall density or the progression to stage IV. This acceleration likely reflects the enhanced amyloidogenic capacity of the G175S PMEL protein, facilitating faster amyloidogenesis and melanosome maturation.

## Discussion

In this study, we present the first near-atomic resolution structures of PMEL amyloids, revealing two distinct polymorphic forms and the structural alterations induced by the G175S mutation. PMEL amyloids play a critical role in melanosomes by providing a scaffold for melanin deposition, which is essential for pigmentation. While PMEL amyloid formation has been studied extensively for its physiological relevance, their high-resolution structure has remained elusive until now ^6, 8, 9, 24^. Our cryo-EM analysis addresses this gap, offering crucial insights into PMEL amyloid architecture and the structural consequences of the G175S mutation associated with PDS.

### Polymorphism and Structural Insights into PMEL Amyloids

Our analysis of native PMEL amyloids revealed two polymorphic forms, Polymorph 1 and Polymorph 2, both adopting a two-protofilament helical architecture. Despite their similar β-sheet configurations, the main chain morphology differs significantly. Polymorph 2 exhibits an inner cavity, absent in Polymorph 1, resulting in a more loosely packed structure. Such polymorphism highlights the structural plasticity of PMEL amyloids, which may facilitate interactions with melanin precursors or melanosomal proteins under varying physiological conditions. This flexibility is a common feature of functional amyloids and may be critical for the regulation of melanosomal architecture and pigmentation ^25–27^.

### Structural Implications of the G175S Mutation

The G175S mutation in PMEL, strongly associated with PDS, introduces a novel hydrogen bond between Ser175 and Tyr159, altering the β-sheet configuration. The mutation divides the first β-sheet (β1) into three shorter segments while preserving the configuration of β2 and β3. These changes enhance amyloid polymerization, as demonstrated by the four-fold increase in polymerization efficiency *in vitro*. Furthermore, G175S fibrils exhibited faster amyloidogenesis in cells, increasing amyloid secretion by ∼70% compared to wild-type PMEL.

Interestingly, cryo-FIB-SEM tomography revealed no structural differences in melanosomes between G175S and wild-type cells, despite increased amyloid secretion. This suggests that while the G175S mutation accelerates amyloidogenesis, the overall melanosomal architecture remains intact. Enhanced amyloidogenic capacity likely allows a greater proportion of PMEL proteins to convert into amyloids, increasing secretion without disrupting melanosomal structure.

### Impact on Melanosome Maturation

Quantification of melanosomal stages revealed a significant increase in stage III melanosomes and a corresponding decrease in stage II in G175S mutant cells, with no changes in stage IV or overall melanosome density. These findings suggest that G175S accelerates the transition from stage II to stage III melanosomes, likely reflecting enhanced amyloidogenic activity. This accelerated maturation aligns with the structural properties of G175S fibrils and may underlie the increased secretion of melanin granules observed in PDS.

### Functional Implications and Pathophysiology of PDS

PDS is characterized by the release of pigment granules into the anterior chamber of the eye, leading to increased intraocular pressure and an elevated risk of PG. Our findings suggest that the structural changes caused by the G175S mutation enhance amyloid formation, promoting excessive melanin granule secretion. This increased secretion may contribute to pathological pigment dispersion, linking PMEL amyloid dysregulation to PDS.

### Roles of the CAF and RPT Domains

Our analysis focused on the CAF domain as the core structural element of PMEL amyloids. However, previous studies have proposed that the repeat (RPT) domain also contributes to PMEL amyloid formation, particularly through its role in organizing fibrillar sheets ^12^. The RPT domain is heavily glycosylated, and these glycosylations are thought to stabilize the sheet-like structures that facilitate melanin deposition ^14^. While our cryo-EM structures did not reveal RPT-derived amyloids in the fibril core, we cannot exclude the possibility that our extraction methods, which involved deglycosylation and protease digestion, may have selectively degraded RPT-derived amyloids or preferentially preserved CAF-derived fibrils. This limitation highlights the need for complementary approaches, such as *in situ* structural analysis, to fully elucidate the roles of CAF and RPT domains in PMEL amyloid architecture.

### Challenges in In Situ Structural Analysis

Although cryo-electron tomography (cryo-ET) has been successfully used to study amyloids in their native cellular contexts, PMEL amyloids pose unique challenges. Their entangled lamellar organization within melanosomes complicates the isolation and visualization of individual fibrils (Figure S6). Furthermore, our 2D classification revealed higher-order assemblies that lack helical twists, suggesting a more complex packing arrangement. These findings indicate that *in situ* structural analysis of PMEL amyloids is technically challenging due to their densely packed and entangled nature.

### Future Directions

Our study provides a foundation for understanding the molecular mechanisms underlying PMEL amyloid formation and its dysregulation in PDS. Future research could focus on exploring the effects of other PMEL mutations, elucidating the role of RPT domain glycosylation, and identifying potential therapeutic strategies to mitigate the pathological impact of accelerated amyloidogenesis in PDS.

In conclusion, our cryo-EM analysis reveals the structural basis of PMEL amyloid polymorphism and the significant impact of the G175S mutation on amyloidogenesis and melanosome maturation. These findings provide new insights into the functional and pathological roles of PMEL amyloids and their broader implications in pigmentation biology and disease.

## Materials and methods

### Cell culture

Human melanoma cell lines HMV-II (TKG-0318) and MNT-1 (CRL-3450) were obtained from the Cell Resource Center for Biomedical Research at Tohoku University and the American Type Culture Collection, respectively. HMV-11 cells were maintained in F12 medium (Fujifilm, Osaka, Japan) supplemented with 10% fetal bovine serum (FBS). MNT-1 cells were maintained in DMEM medium (Fujifilm) supplemented with 20% FBS, 10% AIM-V (ThermoFisher Scientific, Waltham, MA), 0.1 mM MEM non-essential amino acids (Fujifilm).

For isolating native PMEL fibrils, we used the HMV-II human melanoma cell line. MNT1 cells, while also suitable for PMEL expression, were not used for fibril isolation because their melanin granules are too tightly packed to be efficiently disintegrated during extraction. Conversely, MNT1 cells were used for the rest of the experiments. HMV-II cells, although useful for fibril isolation, become unstable after extended culture and multiple passages, leading to a gradual decrease in their capacity to produce melanin granules. Thus, MNT1 cells were more suitable for experiments requiring stable, long-term granule production.

### PMEL gene knockout and transfection

First, we constructed two lentiviral transfer plasmids: pCW57.1-X330-gRNA/SpCas9-mCeru-PuroR and pCW57.1-PMEL-DsRed-HygR-GFP-rTetR (Figure S5a).

### Construction of pCW57.1-X330-gRNA/SpCas9-mCeru-PuroR

The expression cassette backbone for Cas9 and the gRNA scaffold was derived from the pX330_sgRNA/hSpCas9 plasmid (Addgene number 172832) ^28^ . The original gRNA scaffold sequences were replaced with those targeting the PMEL gene.

The sequences of the sgRNA target sites are listed below, with PAM sites underlined:

- AAGTGACTGTCTACCATCGC **<u>CGG</u>**
- CGTGTCCCAGTTGCGGGCCT **<u>TGG</u>**
- TCCATCCAAGGCCCGCAACT **<u>GGG</u>**
- CTCCATCCAAGGCCCGCAAC **<u>TGG</u>**

Following the SpCas9 sequence, we inserted mCerulean3 (synthesized by ThermoFisher Scientific), a T2A sequence, and the puromycin-resistance gene (derived from pCW57.1, Addgene number 99283)^29^. This plasmid enables knockout of the PMEL gene and selection of clones via mCerulean fluorescence and puromycin resistance. The entire cassette, from gRNA to SpCas9 and mCerulean-PuroR, was inserted into pCW57.1 to convert it into a lentiviral transfer plasmid.

### Construction of pCW57.1-PMEL-DsRed-HygR-GFP-rTetR

The backbone for this plasmid is the Tet-on plasmid pCW57.1^30^. We inserted the PMEL gene with or without G175S mutation, along with DsRed (excised from the tdTomato sequence, derived from pCDH-EF1-Luc2-P2A-tdTomato, Addgene number 72486, a gift from Kazuhiro Oka), under the tight TRE promoter. In addition to introducing the G175S mutation, the PAM sites in the PMEL expression plasmid were mutated to prevent cleavage by Cas9. Following the hPGK promoter, we inserted the hygromycin-resistance gene (derived from pCEP4-AD8gp160, Addgene number 123260) ^31^, a T2A sequence, the EGFP gene, another T2A sequence, and the rTetR gene. This plasmid enables doxycycline (Dox)-dependent expression of PMEL G175S and allows clone selection via hygromycin resistance and EGFP fluorescence.

### Virus Production

We transfected Lenti-X 293T cells (Takara Bio, Shiga, Japan) with the transfer plasmids, alongside the packaging plasmid pCMV-dR8.2 delta-vpr (Addgene number 8455) ^32^ and the envelope plasmid pCMV-VSV-G (Addgene number 8454), using the Avalanche-Everyday transfection reagent (EZ Biosystems, College Park, MD). Twenty-four hours post-transfection, the cells were washed three times with PBS and incubated in fresh medium. Virus-containing medium was collected at 48, 72, and 96 hours post-transfection. Cell debris was removed by centrifugation at 1,000×g, followed by filtration through a 0.45 µm membrane (Millipore). The lentivirus-containing supernatant was concentrated using polyethylene glycol precipitation ^33^.

### Generation of PMEL Knockout Cell Lines

To generate PMEL knockout cell lines, HMV-II and MNT1 cells were incubated with the concentrated lentiviral particles carrying pCW57.1-X330-gRNA/SpCas9-mCeru-PuroR for 24 hours. Cells were then selected using 2 µg/ml puromycin. PMEL gene knockout was confirmed via western blotting using a PMEL-specific antibody (E-7, Santa Cruz Biotechnology, Dallas, TX) (Figure S5b). PMEL-knockout cells were subsequently transduced with the concentrated lentivirus carrying pCW57.1-PMEL-DsRed-HygR-GFP-rTetR and selected using 200 µg/ml hygromycin. Expression levels of PMEL were adjusted to match wild-type levels by optimizing Dox concentration (Figure S5b).

We initially attempted to generate PMEL(G175S) expression cells by directly modifying the genomic sequence of the native PMEL gene using CRISPR-Cas9. However, no clones proliferated, likely due to the toxicity of the mutant protein. Consequently, we employed the Tet-on system to control the expression of PMEL(G175S), allowing for Dox-inducible expression and avoiding the issues associated with constitutive expression of the mutant protein.

### Extraction of PMEL amyloids

PMEL amyloid fibrils were isolated following a modified version of the previously described method^8^. Melanoma cells were resuspended in PBS supplemented with 2.5 µg/ml cytochalasin D (Fujifilm) and 10 µM nocodazole (Fujifilm) and incubated at 37℃ for 30 minutes. The cells were then resuspended in 10 mM Tris-HCl (pH 7.4) containing a protease inhibitor cocktail (Nacalai Tesque, Kyoto, Japan) and incubated on ice for 10 minutes.

The cells were disrupted by Dounce homogenization, followed by centrifugation at 800 ×g for 10 minutes at 4℃ to remove cell debris. The membrane fraction was collected by ultracentrifugation at 100,000 ×g for 60 minutes at 4℃, and the resulting pellets were rinsed twice with PBS. Rinsed membranes were lysed in 2% Triton X-100 in PBS for 1 hour at 4℃ and then layered onto a discontinuous sucrose gradient consisting of 30%, 45%, and 55% sucrose in 50 mM Tris-HCl (pH 7.4), 200 mM NaCl, and 1 mM EDTA. The samples were centrifuged for 2 hours at 100,000 ×g at 4℃.

The 55% sucrose fraction was diluted 10-fold and ultracentrifuged at 100,000 ×g for 2 hours at 4℃. The resulting pellet was rinsed twice with PBS and resuspended in 50 mM Tris-HCl (pH 7.4) containing 5 mM CaCl₂ and disrupted by sonication for 10 seconds using a Q125 sonicator (Qsonica, Newtown, CT). For the G175S specimen, the pellet was incubated in 4M Urea in 50 mM Tris-HCl (pH 7.4) for 10 min at room temperature to loosen the structure before sonication. Deglycosylation was carried out by adding 10 µl of O-glycosidase (New England Biolabs, Ipswich, MA) and 10 µl of neuraminidase (New England Biolabs) to the sample, followed by overnight incubation at 37℃.

The digested product was centrifuged at 20,000 ×g for 10 minutes at 4℃, and the pellet was resuspended in 50 mM Tris-HCl (pH 8.0) containing 5 mM CaCl₂ and 1 mM DTT. The pellet was then digested with 10 µg of trypsin for 1 hour at 37℃ ^24, 34^. After centrifugation at 20,000 ×g for 10 minutes at 4℃, the pellet was resuspended in 50 mM Tris-HCl (pH 8.0) containing 5 mM CaCl₂ and 1 mM DTT, followed by digestion with 20 µg of Arg-C endopeptidase overnight at 37℃.

The digested product was centrifuged at 20,000 ×g for 10 minutes at 4℃, then disrupted by sonication for 20 seconds. After ultracentrifugation at 100,000 ×g for 2 hours at 4℃, the pellet was sonicated for an additional 20 seconds. The final specimen was centrifuged at 20,000 ×g for 10 minutes at 4℃ and prepared for cryo-EM analysis (Figure S6).

### In vitro polymerization of PMEL CAF domain

The CAF domain (residues 148-223) of human PMEL was subcloned into the pET24a plasmid (Merck Millipore, Darmstadt, Germany) and transformed into *E. coli* BL21 (DE3) (New England Biolabs). Transformed bacteria were cultured in LB medium at 37℃ until the optical density (OD600) reached 0.8. Expression of the CAF domain was induced by the addition of 0.5 mM isopropyl β-D-thiogalactopyranoside (IPTG), followed by overnight incubation at 18℃. The cells were harvested and disrupted by sonication.

Inclusion bodies were collected and sequentially washed with 2M, 3M, 4M, 6M, and 8M urea. The resulting pellets were solubilized in 6M guanidine hydrochloride (GuHCl), and insoluble debris were removed by centrifugation at 20,000 ×g for 10 minutes at 30℃. The clarified supernatant was loaded onto Ni-NTA resin (Nacalai Tesque) pre-equilibrated with 50 mM Tris-HCl (pH 8.0), 8M urea, and 20 mM imidazole.

The resin was washed with the equilibration buffer, and bound proteins were eluted with 50 mM Tris-HCl (pH 8.0), 8M urea, and 300 mM imidazole. The eluted protein was then diluted with 150 mM sodium acetate (pH 4.4) to reach a final concentration of 0.3 mg/ml, and incubated with vigorous shaking (200 rpm) at 37℃ for 24∼30h for wild-type protein. Initially, we incubated the G175S mutant protein at 37°C; however, the rapid growth of amyloids at this temperature led to the formation of thick, bundled rods that were unsuitable for analysis. To obtain analyzable fibrils, we incubated the G175S protein at 18°C for 30 hours. For the wild-type protein, we incubated at 18°C for 66 hours, resulting in fibrils that exhibited the same structure as those incubated at 37°C.

### Cryo-electron microscopy of PMEL amyloids

Samples were resuspended in 150 mM sodium acetate (pH 4.4) at a final concentration of 0.3 mg/ml. A 2 µl aliquot of the sample was applied to each side of freshly glow-discharged Ultra Au foil R1.2/1.3 300 mesh grids (Quantifoil Micro Tools GmbH, Großlöbichau, Germany). The grids were blotted from both sides for 3 seconds at 12°C under 100% humidity and subsequently plunge-frozen in liquid ethane using a Vitrobot Mark IV (Thermo Fisher Scientific).

Images were recorded on a CRYO ARM 300 II (JEOL, Tokyo, Japan) at the University of Tokyo, operated at 300 keV. An Omega filter with a slit width of 20 eV and a Gatan K3 direct electron detector in correlated-double sampling (CDS) mode were used for imaging. The nominal magnification was set to 60,000×, yielding a physical pixel size of 0.8784 Å/pixel. Movies were acquired using the SerialEM software, with a target defocus range of 0.8-1.5 µm. Each movie was recorded for 5.52 seconds with a total electron dose of 50 e⁻/Å², divided into 50 frames.

### Data processing

Image processing was carried out using CryoSPARC v4.5.3 and Relion v3.1 and v5 ^35, 36^. Raw movie frames were motion-corrected, and the contrast transfer function (CTF) was estimated using the patch-CTF estimation method. Fibrils were automatically picked using the filament tracer tool, and segments were extracted with a box size of 300 pixels and an inter-box distance set to 10% of the box size. Several rounds of 2D classification were performed to exclude bad classes and those that lacked discernible helical twists.

For the native specimen, we identified both 1-start and 2-start helical fibrils (Figure S1, squares), but only the 2-start helical fibrils were processed further. The 1-start helical particles were discarded due to insufficient resolution. In the native wild-type specimen, two distinct polymorphs were identified and processed separately. After selection, the final segments were re-extracted with a box size of 280 (native fibrils) or 300 (*in vitro* fibrils) pixels and reconstructed using helical refinement.

Initial references were generated from 2D class averages with a box size of 600 pixels, using the relion_helix_inimodel2D subroutine ^37^. Further refinement included local and global CTF refinement, reference-based motion correction, and 3D classification. Final maps were generated from 3D classes with the highest resolution using helical refinement. The handedness of the fibrils was determined by generating mirror structures of the 3D maps. Models were built using ModelAngelo ^38^ for both the original and mirrored structures. Only the left-handed structure produced reasonable and consistent models, confirming the fibril handedness.

For the *in vitro* polymerized specimen, additional refinement steps were performed, including Relion CTF refinement and Bayesian polishing after 3D classification. Resolution estimates were determined using the Fourier shell correlation (FSC) at a threshold of 0.143. Image processing workflows were summarized in Figure S1-4.

Initial models were generated using Relion v5.0 ModelAngelo and refined using ChimeraX and ISOLDE tool ^39–41^. The refined models were validated using Phenix (v.1.19.2-4158) ^42^ (Table 1). Image rendering was conducted using Chimera X and MaskChains tool ^43^. We used pyem tools for parameter conversion from CryoSPARC to Relion ^44^.

**Table 1.** Summary of parameters.

|  |  |
| --- | --- |
| Data collection parameters |  |
| **Magnification** | 60,000 |
| **Pixel size (Å)** | 0.8784 |
| **Voltage (keV)** | 300 |
| **Defocus range (µm)** | 0.8-2.0 |
| **Exposure time (sec)** | 5.5 |
| **Number of frames per movie** | 50 |
| **Total dose (e<sup>-</sup>/Å<sup>2</sup>)** | 50 |

|  | Native WT<br>Polymorph 1 | Native WT<br>Polymorph 2 | Native G175S | *In vitro* WT | *In vitro* G175S |
| --- | --- | --- | --- | --- | --- |
| **Number of movies** | 16,081 | 16,081 | 7,280 | 4,331 | 5,020 |
| **Box size (pixels)** | 280 | 280 | 280 | 300 | 300 |
| **Number of particles** | 286,491 | 352,878 | 576,997 | 147,735 | 781,124 |
| **Initial model** | ModelAngelo | ModelAngelo | ModelAngelo | ModelAngelo | ModelAngelo |
| **Bonds lengths**<br>**RSMD (Å)** | 0.090 | 0.089 | 0.090 | 0.089 | 0.088 |
| **Bonds angles**<br>**RSMD (°)** | 1.882 | 1.876 | 1.884 | 1.880 | 1.985 |
| **MolProbability**<br>**score** | 0.91 | 0.70 | 0.69 | 0.91 | 0.91 |
| **Clash score** | 0.00 | 0.00 | 0.00 | 0.00 | 0.00 |
| **Rotamer outliers**<br>**(%)** | 0.00 | 0.00 | 0.00 | 0.00 | 0.00 |
| **Ramachandran**<br>**plot (%)** | Favored: 93.94<br>Allowed: 6.06<br>Outliers: 0.00 | Favored: 96.77<br>Allowed: 3.23<br>Outliers: 0.00 | Favored: 96.88<br>Allowed: 3.12<br>Outliers: 0.00 | Favored: 93.94<br>Allowed: 6.06<br>Outliers: 0.00 | Favored: 93.75<br>Allowed: 6.25<br>Outliers: 0.00 |
| **CaBLAM**<br>**outliers (%)** | 6.45 | 0.00 | 0.00 | 6.45 | 6.67 |
| **B-factors**<br>**(min/max/mean)** | 18.17/55.61/30.58 | 63.33/100.00/98.61 | 80.76/100.00/98.81 | 18.17/55.61/30.58 | 80.76/100.0/98.81 |
| **Map resolution** | 1.79 | 1.79 | 1.79 | 1.94 | 1.79 |
| **estimates (Å)** |  |  |  |  |  |
| **Helical Twist (°)** | 177.7 | -4.1 | 178.4 | 177.8 | 178.4 |
| **Helical Rise (Å)** | 2.34 | 4.67 | 2.35 | 2.31 | 2.38 |
| **EMDB accession numbers** | 61782 | 61783 | 61784 | 61785 | 61786 |
| **PDB accession codes** | 9JST | 9JSU | 9JSV | 9JSW | 9JSX |

### Preparation of specimen for cryo-FIB-SEM

A formvar-coated gold grid stabilized with an evaporated carbon film (FCF100-Au-EC, Electron Microscopy Sciences, Hatfield, PA) was prepared by coating with a 0.1% gelatin solution (Nacalai Tesque) for 1 hour at 37°C. The grid was then washed with culture medium to remove excess gelatin. MNT1 cells were applied to the prepared grid surface and cultured for 96 hours in the presence of 0.2 µg/mL Dox to induce expression of PMEL. Hoechst-33258 was added to the medium at a final concentration of 1 µg/mL and incubated with the cells for 10 minutes to stain nuclei.

Following staining, grids were transferred into a TBS buffer (10 mM Tris-HCl, pH 7.4, 150 mM NaCl) supplemented with 0.1% BSA, 5 mM CaCl₂, 5 mM MgCl₂, and 9% propylene glycol ^44^. The grids were incubated in this buffer for 5 minutes at room temperature to enhance cryoprotection. After incubation, grids were blotted from both sides for 25 seconds at 25°C in a controlled 100% humidity environment using a Vitrobot Mark IV (Thermo Fisher Scientific). Finally, the grids were plunge-frozen in liquid ethane cooled by liquid nitrogen.

### Cryo-FIB-SEM Lamella Preparation

We used an Aquilos cryo-focused ion beam scanning electron microscope (ThermoFisher Scientific) for preparing lamellae^45^ (Figure S7a). The preparation was performed in a stepwise manner, beginning with a lamella thickness of 3 µm, followed by successive milling to reduce the thickness to 2 µm and finally to 0.88 µm. These steps utilized gallium ion beam milling currents of 1 nA, 0.5 nA, and 0.3 nA, respectively. For the final polishing step, milling currents between 13 and 50 pA were applied to reach a final lamella thickness of less than 300 nm. After achieving the desired thickness, the lamellae were coated with an additional thin layer of inorganic platinum by sputter coating at 30 mA for 3 seconds to enhance conductivity and protect the sample during subsequent imaging.

### Cryo-tomography Data Acquisition and Processing

Tomographic tilt series were recorded on a Talos Arctica transmission electron microscope (Thermo Fisher Scientific) operating at 200 keV at the University of Tokyo (Figure S7b). The microscope was equipped with a Gatan Quantum-LS Energy Filter set to a slit width of 30 eV and a Gatan K2 BioQuantum direct electron detector in electron counting mode. Imaging was performed at a nominal magnification of 49,000×, resulting in a physical pixel size of 2.2 Å/pixel. Movies were acquired using Thermo Fisher’s Tomography software, with a target defocus set between 4 and 6 µm. Tilt series were collected over an angular range from -62° to +42°, with a 4.0° increment between images and an initial tilt angle of -10°. Each movie was recorded for 2.0 seconds, with a per-frame dose of 2.0 electrons/Å², subdivided into 15 frames to capture high-resolution details. The total accumulated dose for one tilt series was 54 electrons/Å². This dose was carefully optimized to balance the need for sufficient image contrast and resolution while minimizing electron radiation damage to the carbohydrate-rich structures present in the sample ^47, 48^. Raw tilt series were subjected to motion correction using the Alignframes program in IMOD ^49^. Reconstruction of the corrected tilt series was performed with AreTomo2 ^50^ using a simultaneous iterative reconstruction technique (SIRT) method. Tomograms were denoised using Topaz Denoise3D ^51^. 3D modeling of the tomograms was accomplished using the Drawing tools and Interpolator tool in IMOD.

### Thioflavin T assay

For *in vitro* polymerized fibrils, wild-type and G175S mutant CAF domains (30 µM) were polymerized at 37°C with shaking at 200 rpm. At each time point, 5 µl of 1 mM ThT solution was added to 100 µl of the reaction mixture. Fluorescence was measured using a SpectraMax GeminiEM plate reader (Molecular Devices, San Jose, CA) with an excitation wavelength of 444 nm and an emission wavelength of 480 nm.

For secreted amyloids, PMEL-knockout HMV-II cells, transformed with Tet-on-inducible wild-type or G175S PMEL expression plasmids, were cultured in the presence or absence of 0.2 µg/ml Dox for 96 hours. The culture supernatant from 10 cm dishes was recovered and centrifuged at 800 ×g for 5 minutes to remove cell debris. The clarified supernatant was further centrifuged at 20,000 ×g for 15 minutes at 4°C, and the resulting pellets containing melanin granules were resuspended in PBS. After two washes with PBS, the pellets were demembranated by incubating in PBS containing 1% Triton X-100 for 1 hour at 4°C. The demembranated melanin granules were then washed twice with PBS and resuspended in 100 µl PBS. For ThT fluorescence measurement, 10 µl of the granule suspension was added to 200 µl of 50 µM thioflavin T solution in PBS. Fluorescence was measured as described above using the SpectraMax GeminiEM plate reader. To normalize the ThT fluorescence for secreted amyloid content, the cell number of each culture was determined using a hemocytometer. The fluorescence values were adjusted to reflect secreted amyloids from 1 × 10^7^ cells per data point.

### Ultrathin section electron microscopy of melanosomes

After inducing wild-type or G175S mutant PMEL expression for 96 hours, MNT1 cells were collected and fixed with 4% paraformaldehyde and 0.1% glutaraldehyde for 1 hour at 4°C. The samples were post-fixed with 1% osmium tetroxide, followed by staining with 1% uranyl acetate. Dehydration was performed using a graded ethanol series and acetone, and the samples were embedded in Quetol 812 resin (Nissin EM, Tokyo, Japan). Ultrathin sections of 60 nm were prepared using an ULTRACUT microtome (Reichert Leica) and mounted onto Formvar-coated copper grids. Electron microscopy images were acquired using a JEM-2100F microscope (JEOL, Tokyo, Japan) equipped with an F216 CMOS camera (TVIPS GmbH, Gauting, Germany), operated at 200 keV, at the University of Yamanashi. Melanosomes were identified and counted, and their areas were measured using Fiji software 52.

### Statistical Analysis

For statistical comparison of amyloid yield between the wild-type and G175S mutant CAF domains, two analyses were conducted. First, we used the Mann-Whitney U test due to the non-normal distribution of the data. The test was performed as a two-tailed analysis with an exact *p*-value calculation. The results showed that the G175S mutant yielded significantly higher levels of polymerized amyloids compared to the wild-type (*p* = 0.0079). The test statistic, Z = -2.6548, was outside the 95% confidence interval ([-1.96, 1.96]), indicating a statistically significant difference between the two groups. The observed standardized effect size was 0.79, suggesting a large magnitude of difference in amyloid yield between the wild-type and G175S groups.

Second, for the quantification of secreted amyloids, data are presented as the mean ± standard deviation (SD) from independent experiments (N=8). Statistical analysis was performed using one-way analysis of variance (ANOVA) to test for overall differences between groups (native MNT1, WT Dox (-), WT Dox (+), G175S Dox (-), and G175S Dox (+)). A significant difference was found between the groups (*p* = 7.216e-15). To further analyze pairwise differences, Tukey’s HSD (Honestly Significant Difference) post-hoc test was performed. Tukey’s HSD revealed that the difference between WT Dox (+) and G175S Dox (+) was statistically significant (*p* = 3.36e-5), and there was a highly significant difference between G175S Dox (-) and G175S Dox (+) (*p* = 6.635e-12). Statistical analyses were conducted using the Mann Whitney U test calculator and the ANOVA Calculator (https://www.statskingdom.com/).

For the quantification of melanosome stages, the proportions of stage II, III, and IV melanosomes were compared between wild-type PMEL and G175S mutant PMEL expressing cells using the Wilcoxon signed-rank test, as the proportions of the three stages are dependent (their sum equals 100%). Significant differences were observed in the proportion of stage III melanosomes (*p* = 2.575e-7, effect size r = 0.8358) and stage II melanosomes (*p* = 1.568e-8, effect size r = -0.9054). However, no significant difference was detected for stage IV melanosomes (*p* = 0.3607, effect size r = -0.1464). Statistical analyses were conducted using the online Wilcoxon signed-rank test calculator.

## Acknowledgements

We thank Ms. Natsuko Maruyama (University of Yamanashi) for technical assistance. This research is partially supported by Platform Project for Supporting Drug Discovery and Life Science Research (Basis for Supporting Innovative Drug Discovery and Life Science Research (BINDS)) from Japan Agency for Medical Research and Development (AMED) under Grant Number JP24ama121002. This work was supported by the Takeda Science Foundation (to T.O.), the Japan Society for the Promotion of Science (KAKENHI Grant numbers 21H02654, 24H02285 (to T.O.), 21H04762, 21H04762, and 21H05248 (to M.K.)), and the Human Frontier Science Program (Grant number RGP006/2023 to T.O.).

## Author contributions

T.O. conceived and designed the experiments. H.Y. conducted cryo-EM data collection and analysis. H.A. and H. M. conducted cryo-FIB-SEM tomography. T.O. prepared samples and analyzed the tomography and cell biology/biochemistry data. T.O. and M.K. wrote the manuscript.

## Data availability

The maps and the models are available on the EMDB under the accession numbers shown in Table 1.

