## Supplemental Figures for "Cryo-EM of PMEL Amyloids Reveals Pathogenic Mechanism of Pigment Dispersion Syndrome"

### Supplemental Materials

### Supplemental Figures 1-7

**Supplemental figure 1: Summary of the image processing of native wild type fibrils.**

Initial 2D classification revealed three distinct groups of particles: 1-start/thin filaments (red squares), 2-start/thick filaments (blue squares), and higher-order filament bundles (yellow squares).

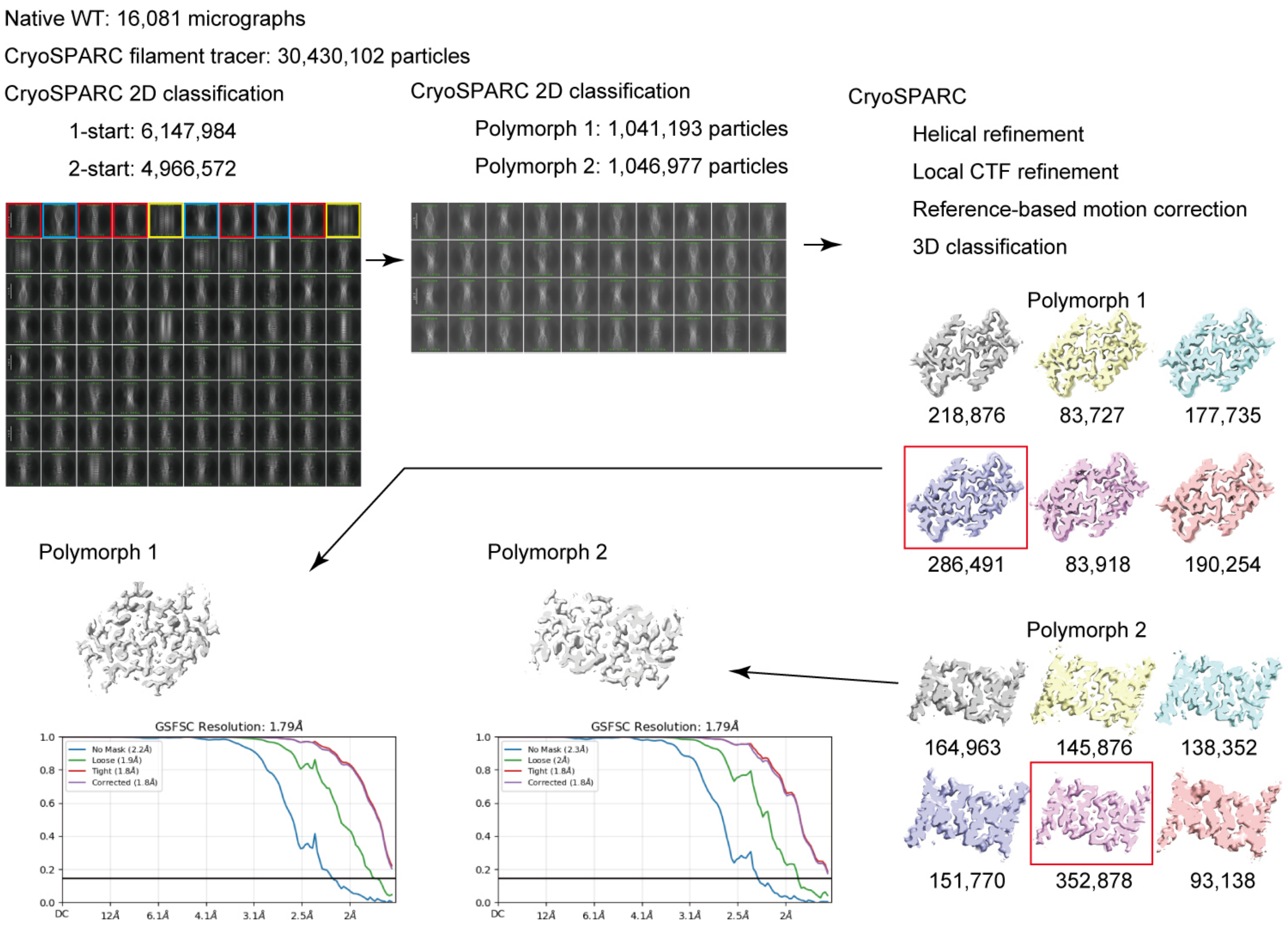

**Supplemental figure 2: Summary of the image processing of native G175S fibrils**

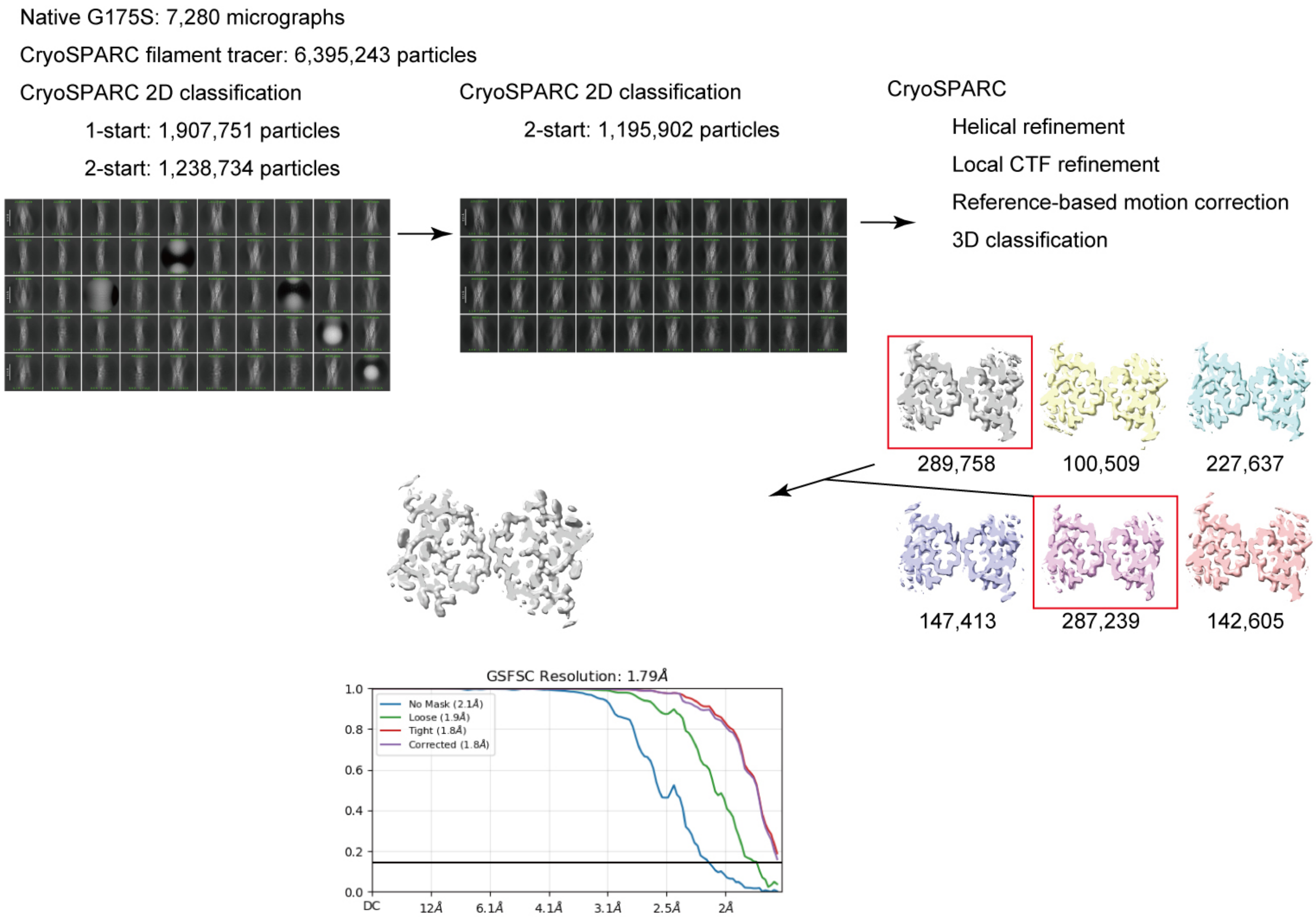

**Supplemental figure 3: Summary of the image processing of in vitro-polymerized wild-type PMEL CAF fibrils**

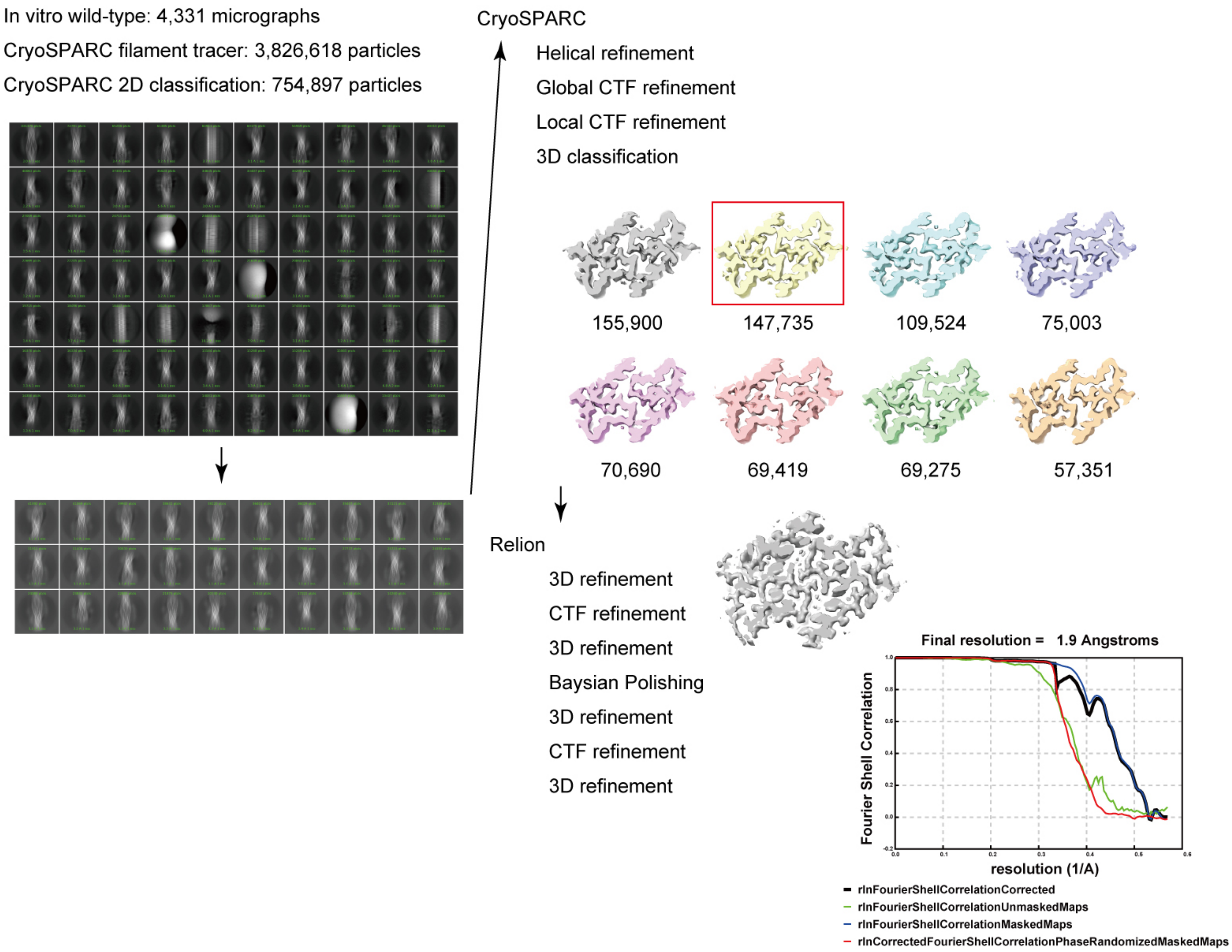

**Supplemental figure 4: Summary of the image processing of in vitro-polymerized G175S PMEL CAF fibrils**

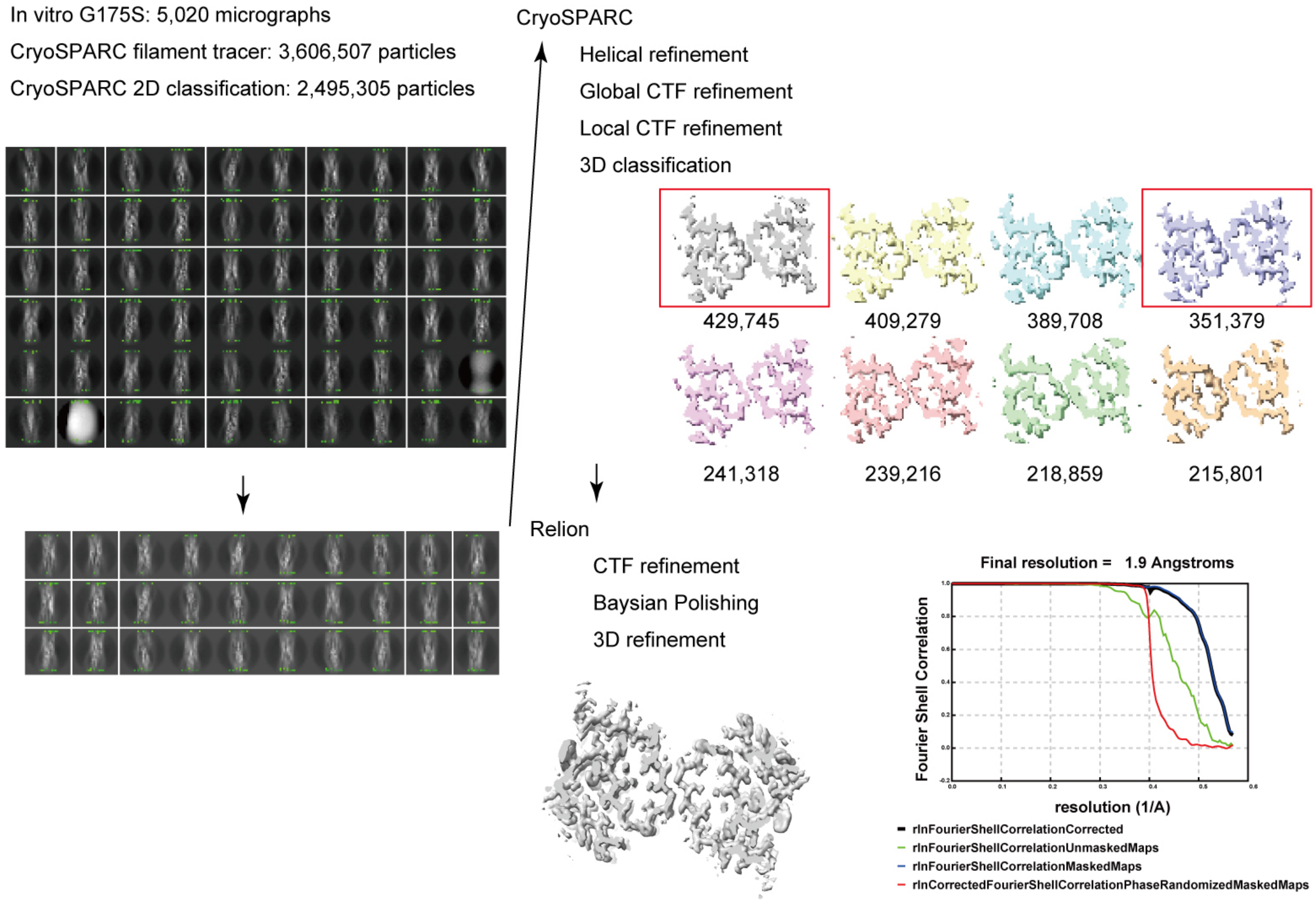

**Supplemental figure 5: Generation of Dox-inducible PMEL WT/G175S Cell Lines**

(a) Vector maps of pCW57.1-X330-gRNA/SpCas9-mCeru-PuroR and pCW57.1-PMEL-DsRed-HygR-GFP-rTetR. The pCW57.1-X330-gRNA/SpCas9-mCeru-PuroR plasmid enables the knockout of the native PMEL gene, while pCW57.1-PMEL-DsRed-HygR-GFP-rTetR allows for doxycycline-inducible expression of the PMEL WT or G175S mutant fused to DsRed.

(b) Western blot analysis of MNT1 cells confirming the knockout of the native PMEL gene (WT/G175S Dox (-)) and the doxycycline-dependent expression of PMEL-DsRed (WT/G175S Dox (+)). Anti-PMEL (E-7) was used to detect both the native PMEL (100 kDa) and the PMEL-DsRed fusion protein (125 kDa). Anti-α-tubulin antibody (B512, Thermo Fisher Scientific) was used as a loading control.

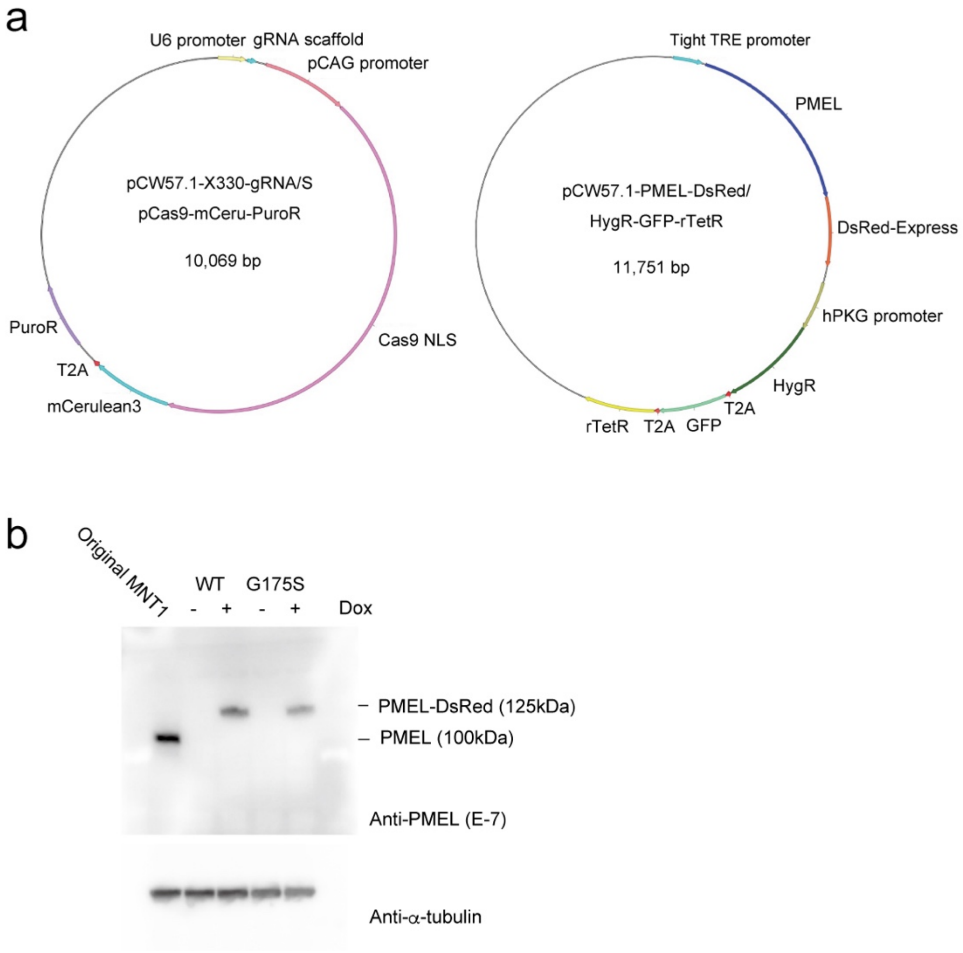

**Supplemental Figure 6: Negative-stain EM images of disintegrated melanin granules for PMEL amyloid extraction.**

Negatively-stained EM images show the fibrillar structures observed after disintegration of melanin granules, highlighting the presence of entangled amyloid fibrils within the lamellar structures.

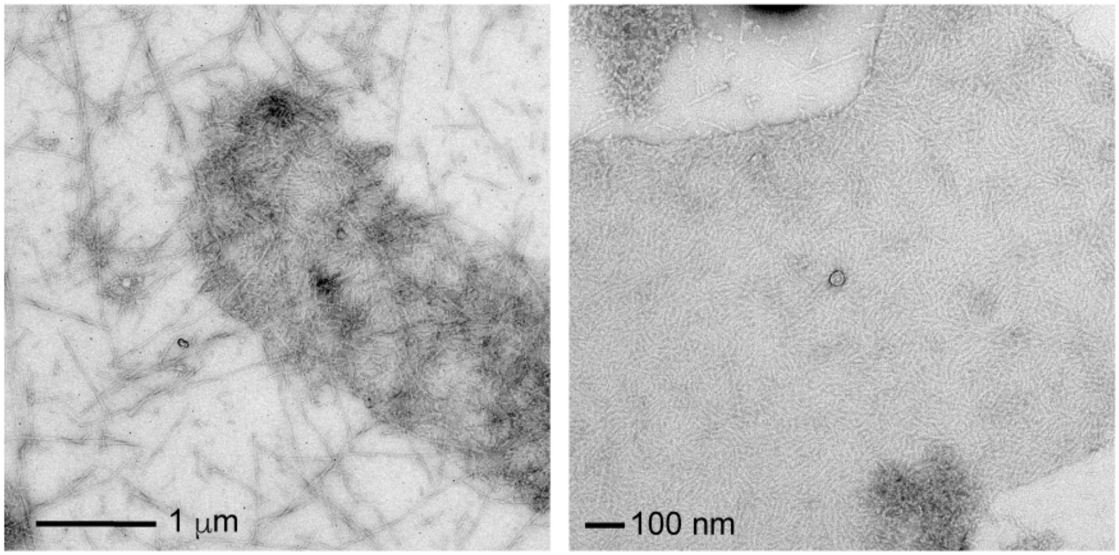

**Supplemental figure 7: Representative images of cryo-FIB-SEM tomography.**

(a) Scanning electron microscopy (SEM) image of an MNT1 cell during lamella targeting, with the targeted area outlined. The dimensions of the patterns are indicated. (b) Transmission electron microscopy (TEM) image of the prepared lamella, showing the dark electron-dense melanosomes.

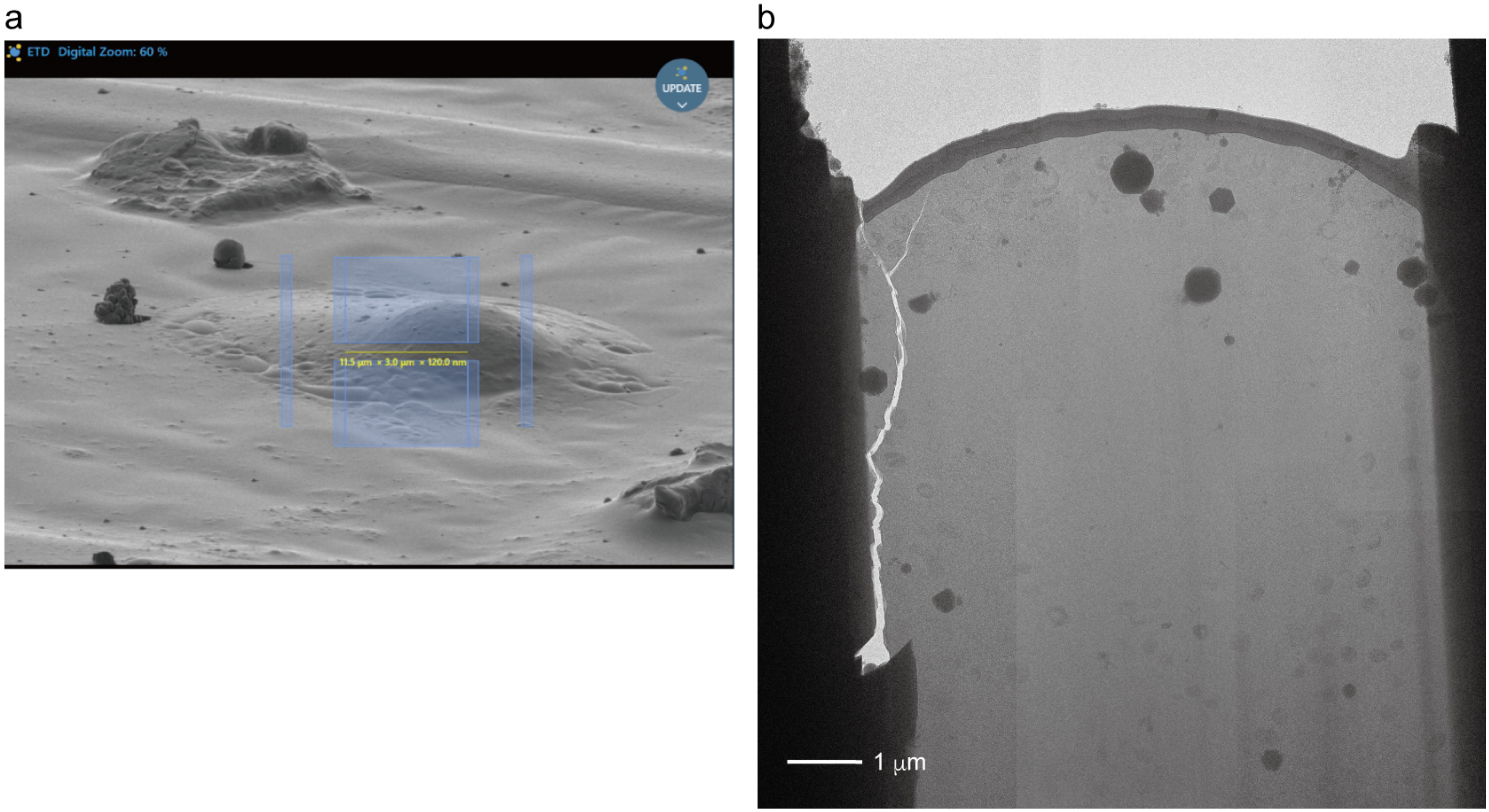
